# A novel mRNA Lipid Nanoparticle Therapy Improves Heart Failure Phenotype and Suppresses Endothelial-Mesenchymal Transition In Vitro

**DOI:** 10.64898/2026.05.22.727265

**Authors:** Muthu Kumar Krishnamoorthi, Sadhna Dhingra, Arvind Bhimaraj

## Abstract

**Objectives:** To evaluate the therapeutic potential of BMP-7 mRNA-lipid nanoparticle formulation in attenuating cardiac fibrosis and improving function in non-ischemic heart failure, and to assess its impact on endothelial phenotype and function under pro endothelial-to-mesenchymal transition (EndMT) conditions.

**Background:** Despite advances in neurohormonal blockade, heart failure (HF) progression remains driven in part by fibrotic remodeling. Endothelial-to-mesenchymal transition (EndMT) has emerged as a contributor to myocardial fibrosis, while recent work suggests endothelial phenotypic plasticity may also participate in myocardial recovery. Bone morphogenetic protein-7 (BMP-7) is a known anti-fibrotic regulator, but effective therapeutic delivery strategies remain limited.

**Methods:** A patent pending, custom-designed BMP-7 mRNA formulated in lipid nanoparticles (AET-1978) was administered subcutaneously in a murine model of non-ischemic HF induced by L-NAME and angiotensin II. Cardiac function and fibrosis were assessed by echocardiography and histology. In an invitro EndMT model, human umbilical vascular endothelial cells (HUVECs) were treated with BMP-7 mRNA and endothelial and mesenchymal morphology, and markers were assessed along with endothelial functional tests.

**Results:** AET-1978 therapy significantly improved left ventricular systolic and diastolic function and reduced myocardial fibrosis compared with untreated HF mice, without evidence of renal or hepatic toxicity. In vitro, BMP-7 mRNA delivery restored endothelial morphology, suppressed EndMT-associated mesenchymal and profibrotic marker expression while preserving nitric oxide production, lipoprotein uptake, and angiogenic capacity of the HUVECs.

**Conclusions:** A novel formulation of BMP-7-mRNA-LNP called AET-1978 represents a novel, transient, non-integrating strategy to attenuate fibrotic remodeling and improve cardiac function in heart failure, with supportive evidence of being anti endothelial to mesenchymal transition.

**Highlights:**

1. A novel BMP-7 mRNA-lipid nanoparticle formulation delivered as a subcutaneous injection attenuated myocardial fibrosis and improved systolic and diastolic function in a murine model of non-ischemic heart failure.
2. BMP-7 mRNA therapy preserved endothelial phenotype and suppressed endothelial-to-mesenchymal transition in an *in vitro* platform of human umbilical vascular endothelial cells.
3. BMP-7 mRNA therapy preserved endothelial function including restoration of nitric oxide production, lipoprotein uptake, and angiogenic capacity *in vitro*.
4. This study introduces AET-1978, a transient, non-integrating mRNA therapeutic platform, as a novel approach to target residual fibrotic pathways in heart failure using a clinically scalable delivery route.

## INTRODUCTION

Pharmacological therapy for heart failure with reduced ejection fraction (HFrEF) has primarily focused on attenuating maladaptive neurohormonal activation through inhibition of the renin–angiotensin–aldosterone system and sympathetic nervous system(1). Although these strategies have substantially improved survival, a significant residual risk of disease progression persists despite optimal medical therapy, highlighting the need to target complementary and incompletely addressed pathological mechanisms(2). Among these, myocardial fibrosis plays a central role in adverse ventricular remodeling and progressive contractile dysfunction(3). While anti-neurohormonal strategy seems to translate into some resolution of fibrosis, a specific therapy that directly targets mechanisms of fibrosis could be complementary.

We have previously identified the role of cell transitions between endothelial and mesenchymal cells in modulating the onset and resolution of fibrosis in a murine model of myocardial recovery from HFrEF(4) and described the presence of endothelial to mesenchymal transitioning (EndMT) cells in end stage human hearts(4). EndMT is characterized by endothelial cells acquiring mesenchymal features under pathological stress, enabling their contribution to fibroblast expansion and extracellular matrix deposition(5). While this mechanism has been implicated in multiple fibrotic diseases, its contribution to cardiac fibrosis and HF has only recently been recognized(6).

Transformation Growth Factor-Beta (TGF-β) is an established strong stimulator of EndMT and previous efforts to block this process using anti-TGF strategy has failed possibly due to the need for a fine balance between fibrotic remodeling for healing vs uncontrolled negative remodeling (7,8). Hence, we chose a strategy that counters the TGF-β pathway to maintain endothelial health and utilizing our mouse model of myocardial recovery and biological reasoning, we identified a target to promote bone morphogenetic protein-7 (BMP-7) as an anti EndMT strategy. BMP-7 is a member of the transforming growth factor-β superfamily with established anti-fibrotic properties (6).

While many genome modifying strategies have been proposed and being studied, a messenger RNA (mRNA) based therapeutic delivered via lipid nanoparticles (LNPs) offers many advantages including a transient, non-integrating approach to restore disease-modifying protein expression with spatial and temporal control(9). Beyond their established use in vaccinology, mRNA-LNP platforms have emerged as a promising modality for cardiovascular applications, yet their potential to modulate fibrotic pathways involving endothelial dysfunction in HF remains incompletely explored (10-12). In this study, we investigated a novel patent pending lipid nanoparticle vehicle loaded with a custom designed mRNA sequence. This particle is being referred to as AET-1978, as a strategy to attenuate fibrotic remodeling in HF.

## METHODS

### *In vivo* murine model

#### Animal model of HFrEF

All animal studies were approved by the Houston Methodist Research Institute Institutional Animal Care and Use Committee and conducted in accordance with the National Institutes of Health Guide for the Care and Use of Laboratory Animals. Twelve-week-old male C57BL/6 mice were randomly assigned to four groups: i) Control, ii) Heart failure with reduced Ejection Fraction (HFrEF), iii) HFrEF + recombinant BMP-7 protein, and iv) HFrEF + AET-1978 **(Figure 1)**. HFrEF was induced using combined L-NAME and angiotensin II (Ang II) administration. Mice in Groups ii–iv received L-NAME (0.3 mg/mL) and 1% NaCl in drinking water, while control mice received normal drinking water. After one week, mice in Groups ii–iv were anesthetized with inhaled isoflurane and implanted subcutaneously with osmotic minipumps (Alzet model 1004) to deliver Ang II at 0.7 mg/kg/day for four weeks. L-NAME and NaCl treatment was continued throughout this period. In the HFrEF + recombinant BMP-7 protein group, a second subcutaneous osmotic minipump was implanted at the time of Ang II pump insertion to deliver recombinant BMP-7 protein for the duration of the study. In the HFrEF + AET-1978 group, subcutaneous injections of AET-1978 was administered weekly beginning at week 1 and continued throughout the study period.

**Figure 1:**
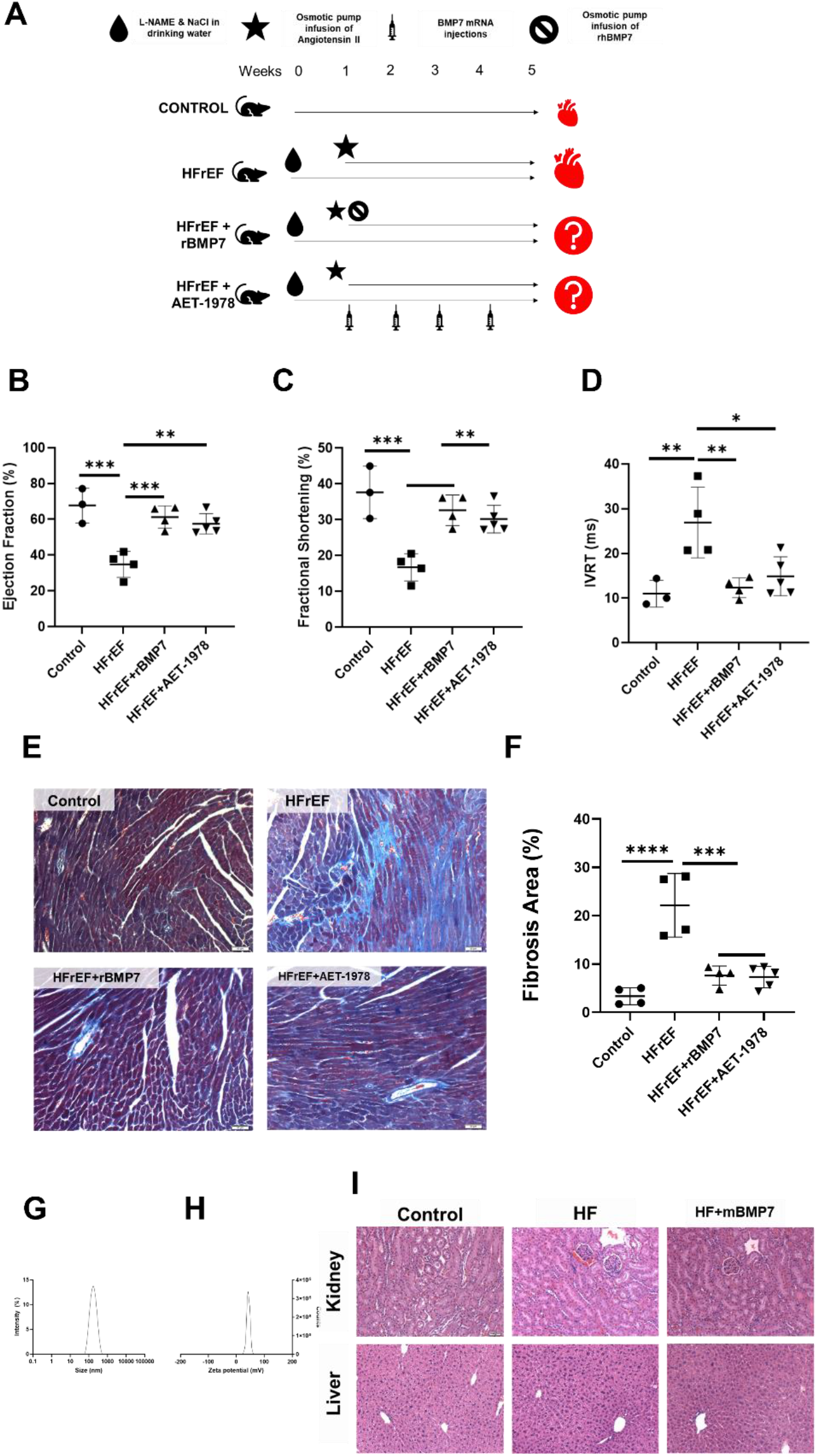
BMP-7 mRNA therapy improves cardiac function and attenuates fibrosis without overt systemic toxicity. (A) Schematic representation of the experimental protocol used to induce non-ischemic heart failure (HF) and administer BMP-7–based therapies.(B) Left ventricular ejection fraction (EF) measured by transthoracic echocardiography at the end of the 5-week protocol.(C) Fractional shortening (FS) assessed by M-mode echocardiography. (D) Isovolumetric relaxation time (IVRT) measured by Doppler echocardiography as an index of diastolic function. (E) Representative Masson’s trichrome–stained transverse heart sections from control, HFrEF, HFrEF + recombinant BMP-7 protein (HFrEF + rBMP7), and HFrEF +BMP-7 mRNA (HFrEF +AET-1978) groups, demonstrating interstitial fibrosis (collagen in blue). Quantification of fibrotic area expressed as a percentage of total myocardial area from Masson’s trichrome–stained sections. (G) Hydrodynamic particle size distribution of BMP-7 mRNA lipid nanoparticles. (H) Zeta potential of BMP-7 mRNA lipid nanoparticles.(i) Representative hematoxylin and eosin (H&E)–stained sections of kidney and liver tissue demonstrating the absence of overt histopathological abnormalities following BMP-7 mRNA treatment. n-3. Data are presented as mean ± SD. Statistical significance was assessed using one-way ANOVA with appropriate post-hoc testing. p < 0.05 was considered statistically significant.

Mice were housed under a 12-h light/dark cycle and fed a standard chow diet (Harlan Teklad 2920). At the conclusion of the five-week protocol, cardiac function was assessed, after which animals were euthanized for tissue collection. Heart, liver, and kidney tissues were harvested at the end of week 5 and processed for histological analysis. Hematoxylin and eosin (H&E) staining was performed on heart, liver, and kidney sections, and Masson’s trichrome staining was performed on heart sections to assess fibrosis. Liver and kidney sections were evaluated by a blinded clinical pathologist and graded for histopathological abnormalities. Heart H&E and trichrome sections were imaged and quantified as previously described (4).

#### Transthoracic echocardiography

Transthoracic echocardiography was performed using a Vevo 2100 imaging system equipped with a 30-MHz MS-400 transducer. Mice were anesthetized with 1–1.5% isoflurane and maintained on a heated platform at 37 °C. Parasternal long- and short-axis views were obtained, and M-mode images were acquired at the level of the papillary muscles from parasternal short-axis views. Left ventricular dimensions and systolic function were quantified using Vevo Lab software (version 5.6). Image acquisition and analysis were performed by separate investigators in a blinded manner.

#### mRNA synthesis

BMP-7 mRNA was generated by the Houston Methodist RNA Core Facility. The human BMP-7 coding sequence (NM_001719) was cloned into a custom, unique, specific optimized mRNA backbone containing 5′ and 3′ untranslated regions and an approximately 150-bp poly(A) tail. In vitro transcription was performed using N1-methyl-pseudouridine-substituted nucleotides to enhance mRNA stability and translational efficiency. Purified mRNA integrity and size were confirmed using an Agilent Tape Station system.

#### AET-1978 synthesis

AET-1978 was formulated with in vivo-jet® lipid nanoparticle reagent according to the manufacturer’s instructions and was administered subcutaneously to the treatment group using standard murine injection method.

#### Physiochemical assessment of lipid nanoparticles

Nanoparticle size and zeta potential were measured by DLS (Malvern Zetasizer). For size, 10 μL of sample was diluted in 990 μL of 1× PBS and loaded into semi-micro disposable cuvettes. For zeta potential, 10 μL of sample was diluted in 90 μL of 1× PBS and 900 μL of ddH_2_O to reduce salt concentration and prevent electrode damage. Measurements were performed in triplicate and averaged to obtain final values.

#### In vitro experiment

Human umbilical vein endothelial cells (HUVECs; Lonza, C2519A) at passages 2–5 was cultured under standard conditions (37 °C, 5% CO_2_) in complete EGM™-2 medium. The medium was washed and replaced with induction agents as described below.

#### Endothelial-to-mesenchymal transition (EndMT) in vitro platform

We utilized a previously described in vitro EndMT model (13) where EndMT is induced by replacing growth medium with EBM™-2 containing 1 mM Nω-nitro-L-arginine methyl ester (L-NAME) and 1 μM angiotensin II (Ang II) on Day 1 and 3 of the experiment. Cells were maintained under EndMT-inducing conditions for 4 days. Where indicated, In vitro transfection of BMP 7 mRNA was performed using the TransIT® mRNA Transfection Kit (MirusBio, USA), following the manufacturer’s protocol concomitantly. At the end of the experiment, cells were fixed with 4% paraformaldehyde for immunofluorescence analysis(13). Expression of endothelial and mesenchymal markers were assessed as described below. Endothelial function assessment (as described below) was performed on Day 4.

Immunofluorescence staining was performed as described in our previous studies (13). Briefly, fixed cells were washed, permeabilized, and blocked prior to incubation with primary antibodies against CD31 (Abcam), VE-Cadherin (Santa Cruz), Vimentin (Santa Cruz), Collagen I (Cell Signaling), and Transgelin (Abcam), each used at a dilution of 1:100. Secondary antibodies were applied at a dilution of 1:200. Nuclei were counterstained with DAPI, and fluorescent images were captured using an EVOS fluorescence microscope. Image analysis was performed using ImageJ (14).

### *In vitro* EndMT functional assays

#### Nitric oxide (NO) production

After 4 days of EndMT induction with or without target therapy, HUVECs cultured in 8-chamber slides were washed three times with Hank’s balanced salt solution (HBSS) containing calcium and magnesium. Cells were incubated with 8 μM DAF-FM™ diacetate in HBSS for 30 min at 37 °C, followed by three washes with HBSS. Fresh EGM™-2 medium was then added, and cells were incubated for an additional 15 min to allow de-esterification. Fluorescent detection of NO was performed using a FITC filter set on a fluorescence microscope.

#### Acetylated Dil-LDL uptake

Following EndMT induction with or without target therapy, HUVECs were washed three times with HBSS and incubated with DiI-acetylated low-density lipoprotein (DiI-AcLDL) working solution for 4 h at 37 °C. Cells were washed again with HBSS, and uptake was visualized using a TRITC filter cube on a fluorescence microscope.

#### *In vitro* angiogenesis (tube formation) assay

After 4 days of EndMT induction with or without target therapy, HUVECs were detached using trypsin-EDTA, neutralized with complete medium, and centrifuged at 200–300 × g for 4 min at room temperature. Cells were resuspended in pre-warmed culture medium, and viable cells were counted using trypan blue exclusion. Growth factor-reduced Matrigel was thawed at 4 °C and aliquoted (140 μL per well) into pre-chilled 48-well plates, which were incubated at 37 °C for 30 min to allow gelation. A total of 30,000 cells in 400 μL medium were then seeded onto the Matrigel surface. Plates were incubated at 37 °C, 5% CO_2_, and tube formation was monitored every hour. Images were acquired using an inverted microscope (×4 or ×10 objective), and quantitative analysis of tube networks was performed using ImageJ.

#### Statistics

Statistical analyses were performed using Prism (version 10; GraphPad, San Diego, CA). Comparisons between two groups were conducted using Student’s t-test. Comparisons among multiple groups were performed using one-way or two-way ANOVA with Tukey’s post hoc correction, as appropriate. A p value <0.05 was considered statistically significant. Data are presented as mean ± SD with individual data points shown.

## RESULTS

Compared to control mice, HFrEF animals exhibited significant left ventricular systolic dysfunction, manifested by reduced ejection fraction and fractional shortening. Treatment with AET-1978 significantly improved systolic function, preserving both ejection fraction and fractional shortening along with preservation of diastolic function as reflected by IVRT measurement when compared to untreated HFrEF mice. The improvement of echo parameters was comparable to that achieved with recombinant BMP-7 protein alone (Fig. 1B–D).

Histological analysis showed marked interstitial fibrosis in HFrEF hearts compared to controls, as demonstrated by Masson’s trichrome staining. AET-1978 treatment substantially attenuated such myocardial fibrosis showing a significantly reduced fibrotic area compared with untreated HFrEF mice (Fig. 1E,F). Physicochemical characterization of the AET-1978 formulation demonstrated nanoparticle size distribution (average particle size - 150 nm) and surface charge (average zeta potential - +50 mV) suitable for in vivo application (Fig. 1 G, H) (15)Blinded histopathological evaluation of kidney and liver tissue revealed no overt differences in abnormalities in the AET-1978 treated animals compared to HFrEF animals, supporting a lack of a systemic impact. (Fig. 1 I).

To provide cellular context for the observed in vivo effects, we examined the impact of BMP-7 mRNA on endothelial phenotype using an in vitro model of endothelial-to-mesenchymal transition (EndMT) in human umbilical vein endothelial cells (HUVECs) (Fig. 2A). EndMT induction resulted in morphological changes consistent with mesenchymal transition, including loss of cobblestone architecture and elongation of cells. Treatment with AET-1978 preserved endothelial morphology under EndMT conditions (Fig. 2B). Successful translation of delivered BMP-7 mRNA using AET-1978 was confirmed by intracellular BMP-7 protein expression detected by immunofluorescence, with corresponding increases in fluorescence intensity (Fig. 2C–D). Consistent with phenotypic preservation, while EndMT induction significantly increased expression of mesenchymal and profibrotic markers, including vimentin, transgelin (SM22α), and collagen I, AET-1978 treatment suppressed the upregulation of all three markers compared with untreated endothelial cells that underwent EndMT (Fig. 3A–F), indicating attenuation of mesenchymal transition.

**Figure 2:**
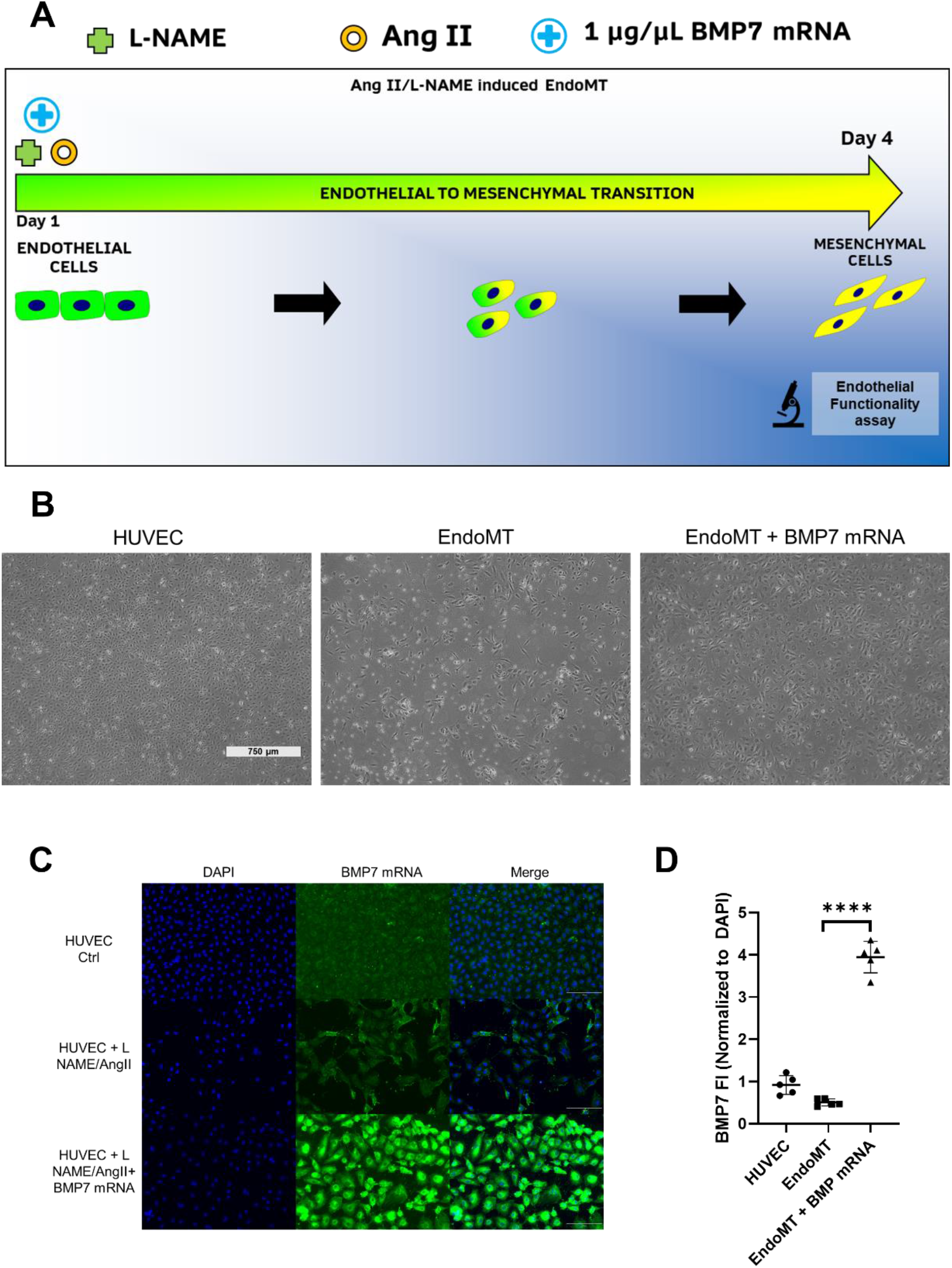
BMP-7 mRNA restores endothelial phenotype and suppresses EndMT in human endothelial cells. (A) Schematic representation of the in vitro endothelial-to-mesenchymal transition (EndMT) experimental design in HUVECs, including EndMT induction and BMP-7 mRNA treatment.(B) Representative phase-contrast images of HUVEC morphology under control conditions, following EndMT induction, and following EndMT induction with BMP-7 mRNA treatment.(C) Representative immunofluorescence images demonstrating intracellular BMP-7 protein expression in HUVECs following BMP-7 mRNA treatment.(D) Quantification of BMP-7 immunofluorescence intensity across experimental groups. n=3. Data are presented as mean ± SD. Statistical significance was assessed using one-way ANOVA with appropriate post-hoc testing. p < 0.05 was considered statistically significant.

**Figure 3:**
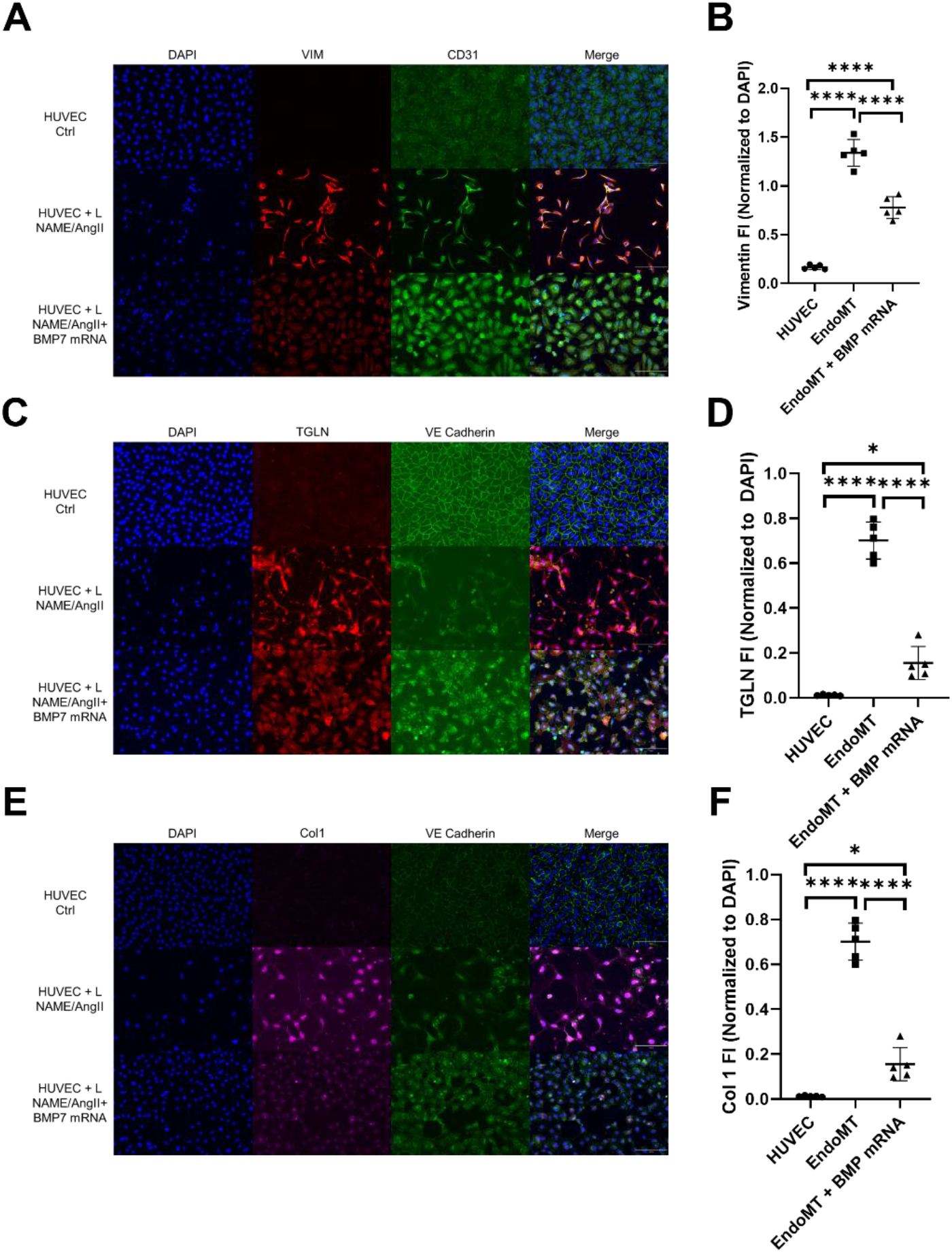
BMP-7 mRNA suppresses mesenchymal and profibrotic marker expression during EndMT. (A) Representative immunofluorescence images showing vimentin expression in HUVECs under control conditions, following EndMT induction, and following EndMT induction with BMP-7 mRNA treatment.(B) Quantification of vimentin fluorescence intensity (FI).(C) Representative immunofluorescence images showing transgelin (SM22α) expression across experimental groups.(D) Quantification of transgelin fluorescence intensity.(E) Representative immunofluorescence images showing collagen I (Col1) expression.(F) Quantification of collagen I fluorescence intensity. Nuclei are counterstained with DAPI. n=3. Data are presented as mean ± SD. Statistical significance was assessed using one-way ANOVA with appropriate post-hoc testing. p < 0.05 was considered statistically significant.

Finally, we assessed whether AET-1978 preserved the endothelial function amongst the cells that it prevented transitioning. EndMT induction resulted in reduced endothelial function reflected by nitric oxide production and impaired acetylated LDL uptake. AET-1978 treatment significantly restored both nitric oxide production and LDL uptake (Fig. 4A–B) proving maintenance of endothelial functionality. EndMT markedly impaired angiogenic capacity of the endothelial cells in Matrigel tube formation assays, whereas AET-1978 treatment restored capillary-like network formation, as evidenced by increased numbers of network segments and greater average segment length (Fig. 4C– E). Together, these findings demonstrate that AET-1978 not only suppresses EndMT-associated mesenchymal marker expression but also preserves core endothelial functions.

**Figure 4:**
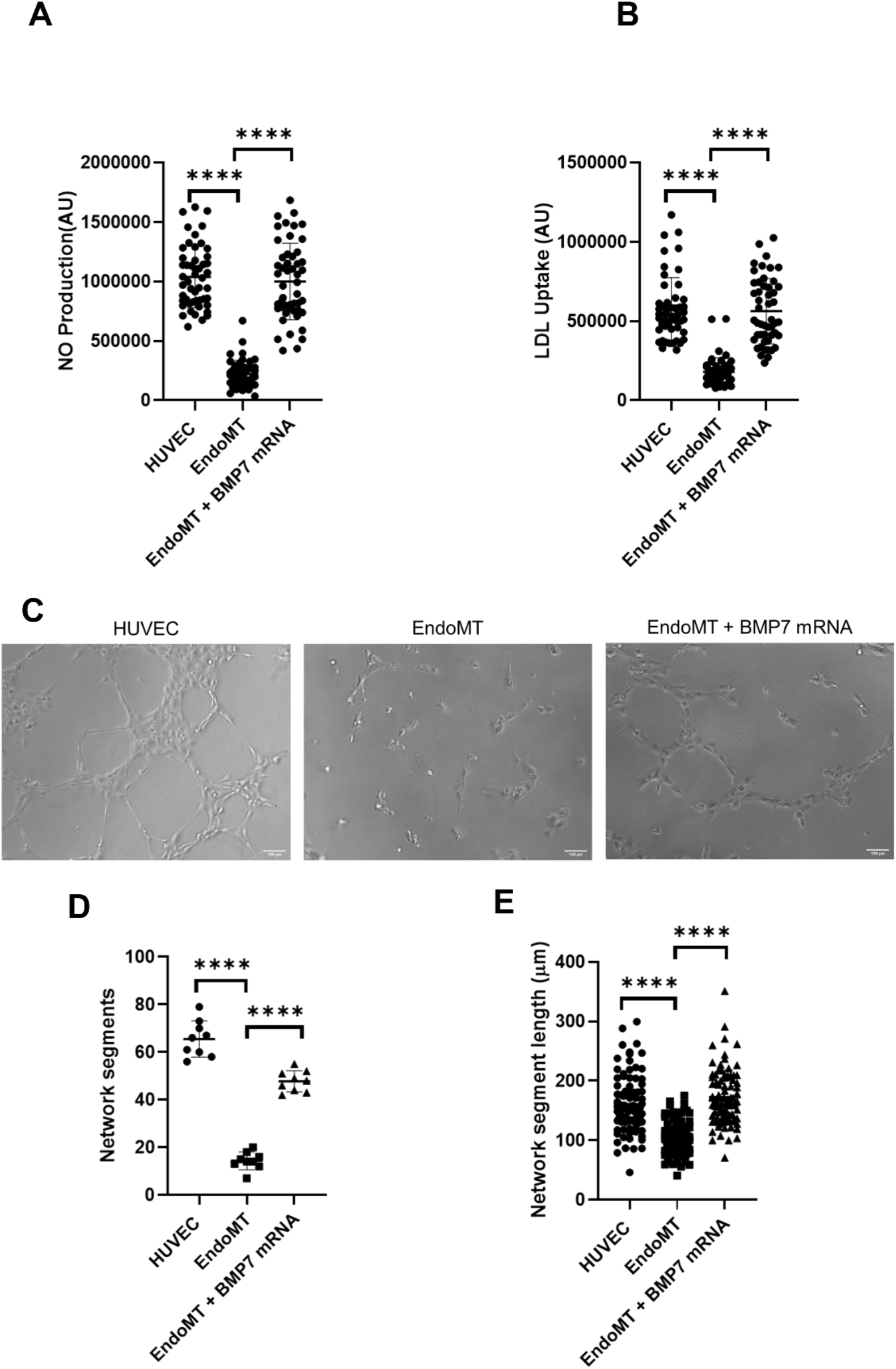
BMP-7 mRNA preserves endothelial function during EndMT. (A) Nitric oxide (NO) production assessed by DAF-FM diacetate fluorescence in control HUVECs, following EndMT induction, and following EndMT induction with BMP-7 mRNA treatment.(B) Uptake of acetylated DiI-LDL in HUVECs under control conditions, EndMT, and EndMT with BMP-7 mRNA treatment.(C) Representative bright-field images of capillary-like tube formation in HUVECs subjected to control conditions, EndMT, or EndMT with BMP-7 mRNA treatment.(D) Quantification of network segments formed during the tube formation assay.(E) Quantification of average network segment length in the tube formation assay. n=3. Data are presented as mean ± SD. Statistical significance was assessed using one-way ANOVA with appropriate post-hoc testing. p < 0.05 was considered statistically significant.

## DISCUSSION

In this study, we demonstrate that a novel, patent pending m-RNA-LNP formulation (AET-1978) targeting BMP-7 and administered via subcutaneous injection, attenuates myocardial fibrosis and improves cardiac function in a murine model of non-ischemic heart failure. This delivery route enables effective clinical translation targeting cardiac therapeutic pathways while remaining compatible with repeat, chronic administration. Endothelial phenotype preservation and suppression of endothelial-to-mesenchymal transition were demonstrated in human endothelial cells exposed to EndMT-inducing conditions, providing mechanistic cellular context for these in vivo findings. Together, these results extend prior work implicating BMP-7 in anti-fibrotic signaling(6,16) by introducing a transient, non-integrating mRNA-based strategy capable of promoting a pro-endothelial health milieu through BMP-7 expression during heart failure. The favorable physicochemical properties of AET-1978 and absence of overt renal or hepatic toxicity further support the translational potential of such a therapy to target residual remodeling mechanisms in heart failure that remain inadequately addressed by current therapies.

Several limitations of this study should be acknowledged. While our in vivo findings establish therapeutic efficacy, the cellular studies presented here were intentionally focused on endothelial-to-mesenchymal transition (EndMT) and endothelial functional integrity, reflecting prior observations from our laboratory and a hypothesis-driven mechanistic focus. As such, the beneficial effects of AET-1978 observed in vivo could be mediated beyond endothelial cells. Furthermore, direct assessment of EndMT or cell-specific mRNA uptake within cardiac tissue was not performed; future studies incorporating lineage tracing, biodistribution analyses, or single-cell profiling will be required to more precisely validate the mechanisms underlying AET-1978–mediated cardio-protection. Finally, the present study evaluated short-term functional and histological outcomes, and long-term durability, repeat dosing effects, and comprehensive toxicological profiling remain to be established.

## CONCLUSION

In conclusion, we demonstrate that lipid nanoparticle–mediated, subcutaneous delivery of a specific construct of BMP-7 mRNA, formulated as AET-1978, improves cardiac function and attenuates myocardial fibrosis in a murine model of non-ischemic heart failure. Using a transient, non-integrating mRNA platform and a clinically relevant delivery route, this approach enables restoration of BMP-7 signaling and engages disease pathways not directly targeted by current neurohormonal therapies. Collectively, these findings support AET-1978 therapy as a promising foundation for next-generation, RNA-based interventions targeting residual fibrotic and remodeling pathways in heart failure.

### Clinical competencies

Our report highlights the limitations in our current medical knowledge of the pathobiology of heart failure and how the prevailing paradigm of therapy for heart failure focusses on cardiomyocytes. The biological variations for therapeutic response seen in the clinic might be a result of specific therapies that target the non-cardiomyocytes.

### Translational outlook

This study highlights a potential therapeutic strategy with a novel delivery mechanism that can target fibrotic remodeling in heart failure (HF). The demonstration that subcutaneous delivery of a BMP-7 mRNA-LNP formulation improves cardiac function and reduces fibrosis suggests a potential complementary approach that may be integrated alongside existing HF treatments. The use of a transient, non-integrating mRNA platform and a minimally invasive delivery route supports clinical feasibility for possible compounding therapy or to partially substitute oral medications.

## Acknowledgement

This work was supported by institutional funds.

